# Interaction of FlhF, SRP-like GTPase with FliF, MS ring component assembling the initial structure of flagella in marine *Vibrio*

**DOI:** 10.1101/2023.01.12.523874

**Authors:** Yuria Fukushima, Michio Homma, Seiji Kojima

## Abstract

*Vibrio alginolyticus* forms a single flagellum at its cell pole. FlhF and FlhG are known to be the main proteins responsible for the polar formation of single flagellum. FlhF, which belongs to the signal recognition particle (SRP)-type GTPase family, localizes at the cell pole and initiates flagellar generation. In contrast, FlhG negatively regulates flagellar numbers. Furthermore, MS-ring formation in the flagellar basal body appears to be an initiation step for flagellar assembly. The MS-ring is formed by a single protein, FliF, which has two transmembrane (TM) segments and a large periplasmic region. We had shown that FlhF was required for the polar localization of *Vibrio* FliF, and FlhF facilitated MS-ring formation when FliF was overexpressed in *E. coli* cells. These results suggest that FlhF interacts with FliF to facilitate MS-ring formation. Here, we attempted to detect this interaction using *Vibrio* FliF fragments fused to a tag of Glutathione S-transferase (GST) in *E. coli*. We found that the N-terminal 108 residues of FliF, including the first TM segment and the periplasmic region, could pull FlhF down. In the first step, the SRP and its receptor are involved in the transport of membrane proteins to target them, which delivers them to the translocon. FlhF may have a similar or enhanced function as SRP, which binds to a region rich in hydrophobic residues.

**IMPORTANCE:** *Vibrio alginolyticus* forms only a single flagellum at the cell pole by regulators of FlhF and FlhG. FlhF regulates positively the formation of flagella and is required for polar positioning of the flagellum. FliF, the two transmembrane (TM) segments and a large periplasmic region, forms the MS ring of flagellar basal body in the membrane. Previous studies suggest that FlhF interacts with FliF to facilitate MS ring formation at the cell pole, but the interaction has not been detected. Here, we show the evidence that FlhF interacts with FliF at residues including the first TM segment and following periplasmic region. The hydrophobic residues of this region seem to be important for the interaction.

## INTRODUCTION

Many motile bacteria possess flagella, whose number and location are precisely controlled among species. *Pseudomonas aeruginosa* and *Vibrio* have a single flagellum on one pole of the cell. *Campylobacter* has a single flagellum on both poles, and *Helicobacter pylori* has multiple flagella on one pole. *Escherichia coli, Bacillus subtilis*, and *Salmonella* have multiple flagella on their entire cell surface. In bacteria, localization on the cell surface is regulated by ATPase or GTPase to assemble structures (1, 2). *Vibrio alginolyticus* has a single flagellum at the cell pole, and its number and location are regulated by FlhF, a GTPase, and FlhG, an ATPase (3). If FlhF is overexpressed in *Vibrio*, it results in the synthesis of multiple polar flagella (4). However, the loss of FlhF results in the absence of a flagellum. FlhF, which is a homolog of a signal recognition particle (SRP) protein and its receptor, is known to be a positive regulator of flagellar number. The SRP and its receptor are involved in the translocation of the Sec system into the membrane through protein targeting (5, 6). Thus, it has been speculated that FlhF may function similarly to SRP. In contrast, FlhG negatively regulates flagellar numbers. Overproduction of FlhG causes a decrease in the number of flagella, whereas depletion of *FlhG* increases the number of flagella. FlhF and FlhG interact with each other (7). Additionally, the reactions of GTP and ATP by FlhF and FlhG, respectively, appear to be involved in the regulation of flagellar numbers. Research has shown that the polar localization of FlhG is dependent on the membrane protein HubP (8), and in the marine *Vibrio* or *V. alginolyticus*, the defect in HubP causes cells to be multiflagellated and to lose the polar localization of FlhG (9). Furthermore, the membrane protein SflA is present at the poles in a HubP-dependent manner (10), and the defect in SflA causes cells to generate peritrichous flagella in the *flhF* and *flhG* double mutant which cannot form the flagella. There is speculation that SflA prevents the formation of multiple flagella other than polar flagella (11). The C-terminal cytoplasmic and N-terminal periplasmic regions of SflA have a DnaJ domain and a tetratricopeptide repeat and Sel1-like repeat motif, respectively (10, 12). We found that mutations of FliM lead to the multiple flagella phenotype and that FliM interacts with FlhG and increases its ATPase activity (13). Furthermore, we showed that FlhG interacts with FlaK, a flagellar master regulator, to suppress flagellar gene expression (14). These factors interact in a complicated manner to regulate the number of flagella.

Flagellar motors at the base of the flagellum rotate the flagellum like a screw as bacteria swim through a liquid. The flagellar motor consists of a stator and rotor, whose interaction generates rotational forces driven by ion motive forces. The MS-ring, located at the flagellar base, is a part of the flagellar rotor and has a complicated structure (15, 16). FliF, which has two transmembrane domains and a large periplasmic region, forms an MS-ring in the cytoplasmic membrane. The MS-ring is one of the earliest structures assembled in the flagellar basal body. Recently, the atomic structure of the MS-ring of *Salmonella* was reported using cryo-EM and its crystal structure (17–19). The MS ring is formed by the assembly of 34 molecules of FliF, and three ring-building motifs (RBM), named RBM1, RBM2, and RBM3 from the N-terminus, have been identified in the periplasmic region of FliF. With 34 molecules, RBM3 forms a ring known as the S-ring. Furthermore, RBM2 forms an M ring with 21–22 molecules. RBM1 creates an M ring with RBM2. RBM1 and RBM2 have different conformations in the FliF molecules, forming an M-ring structure with RBM2. The details of how the periplasmic region of FliF assembles the complex of the MS ring structure have not been clarified. Additionally, we know that the C-ring, one of the rotor structures, is created using three different proteins, FliG, FliM, and FliN, and interacts with the MS ring at the C-terminal region of FliF via FliG (20).

Our previous research suggested that FlhF is required for the polar localization of *Vibrio* FliF and that FlhF facilitates MS-ring formation when FliF is overproduced in *E. coli* cells (21); this evidence suggests that FlhF interacts with FliF to facilitate MS-ring formation at the cell pole; however, the interaction between FlhF and FliF has not yet been directly detected. In this study, we attempted to detect the interaction of FliF with four *Vibrio* FliF fragments fused to Glutathione S-transferase (GST) in *E. coli*.

## RESULTS

### Design and purification of FliF fragments

To investigate the interaction sites of FlhF and FliF, we constructed four FliF fragments and a wild-type FliF (Fig. 1 and Fig. S1): 1) N-terminus (1–55) without the first transmembrane region; 2) N-terminus (1–80) containing the first transmembrane region; 3) periplasmic region (74–469); and 4) C-terminus (493–580) after the second transmembrane region. The fragments were cloned into the vector pGEX-6P-2, and GST was fused to the N-terminus of the fragments, with 11 residues between the GST and FliF containing the PreScission protease recognition site. The resultant plasmids were named pYF101, pYF102, pYF103, and pYF104, which produced GST fusion proteins GST-FliF_1, GST-FliF_2, GST-FliF_3, and GST-FliF_4, respectively. The calculated sizes of GST-FliF_1, GST-FliF_2, GST-FliF_3, and GST-FliF_4 are 32.6 kDa, 35.5 kDa, 70.8 kDa, and 36.3 kDa, respectively. The proteins were well expressed and detected at the corresponding molecular weights, except for pYF105, which produces wild-type FliF (Fig. S2). At 37°C, most of the FliF_2 was precipitated as a pellet when it was expressed, and the cytoplasmic fraction of disrupted cells was centrifuged. Thus, after cells reached 0.5 of OD_660_ at 37°C, GST-FliF_2 was induced by adding IPTG and incubating cells at 18°C for approximately 3 h. This treatment allowed GST-FliF_2 to fractionate into a soluble cytoplasmic fraction. The soluble fractions were then loaded into 1 ml of GSTrap (FF) (GST column), the column was washed with PBS, and the protein was eluted five times (1 ml of the elution buffer). We detected proteins mostly in elutions one and two; thus, these fractions were passed through a desalting column. The protein concentrations are 0.37 mg/ml, 0.13 mg/ml, 0.13 mg/ml, and 0.32 mg/ml for FliF_1, FliF_2, FliF_3, and FliF_4, respectively (Fig. 2). After desalting, aliquots of the solutions were frozen in liquid nitrogen and stored at −80°C until use.

**FIG 1.**
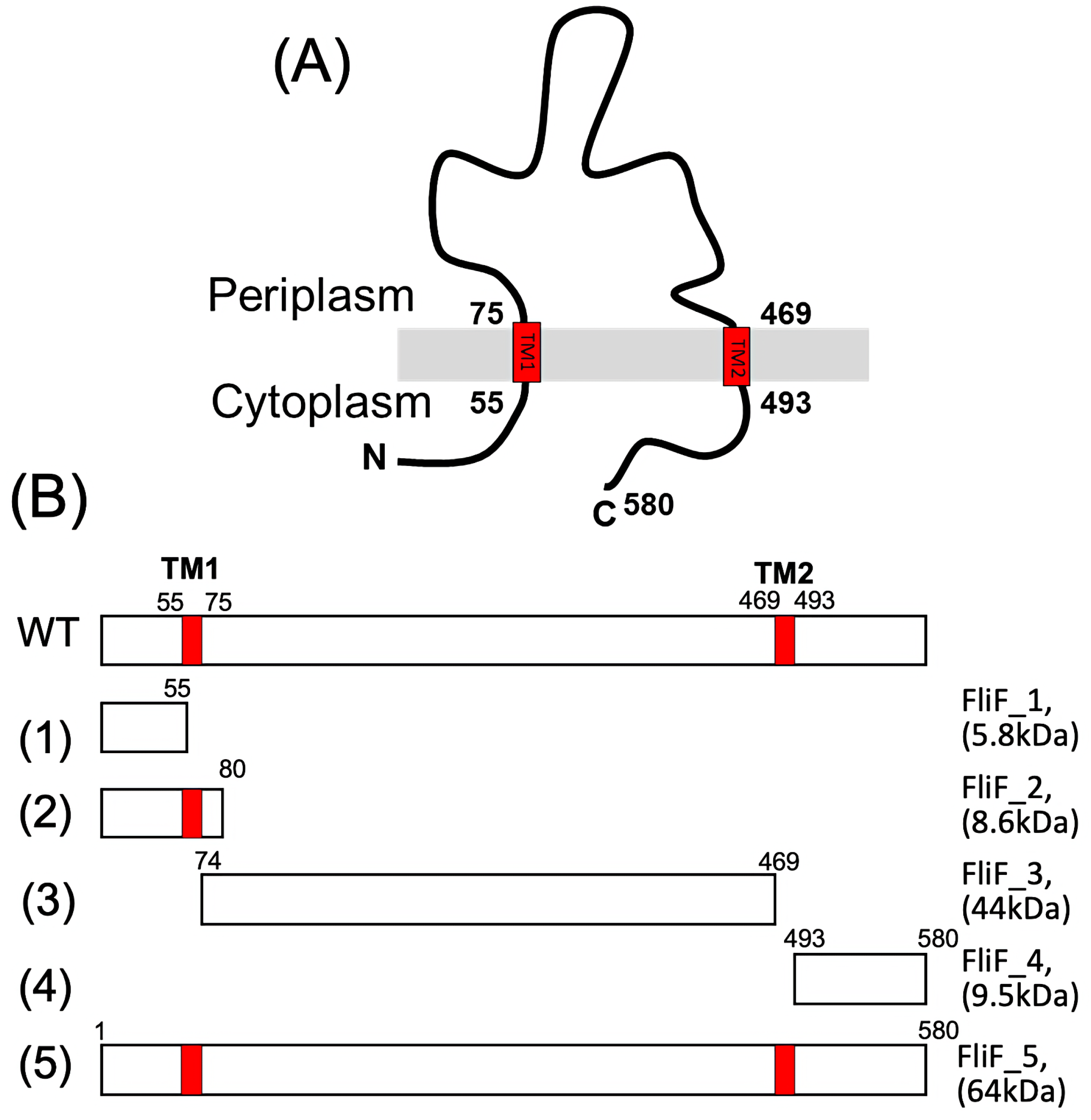
(A) The membrane topology of a *Vibrio* FliF protein. The red rectangles in the membrane (gray bar) show the transmembrane regions of FliF which has two transmembrane (TM) regions. (B) Schematic primary structure of the FliF fragment constructed in this study. The fragments are produced from plasmids (1) pYF101, (2) pYF102, (3) pYF103, (4) pYF104, and (5) pYF105 and named FliF_1, FliF_2, FliF_3, FliF_4, and FliF_5. The expected molecular weights of the fragments are shown in parentheses.

**FIG 2.**
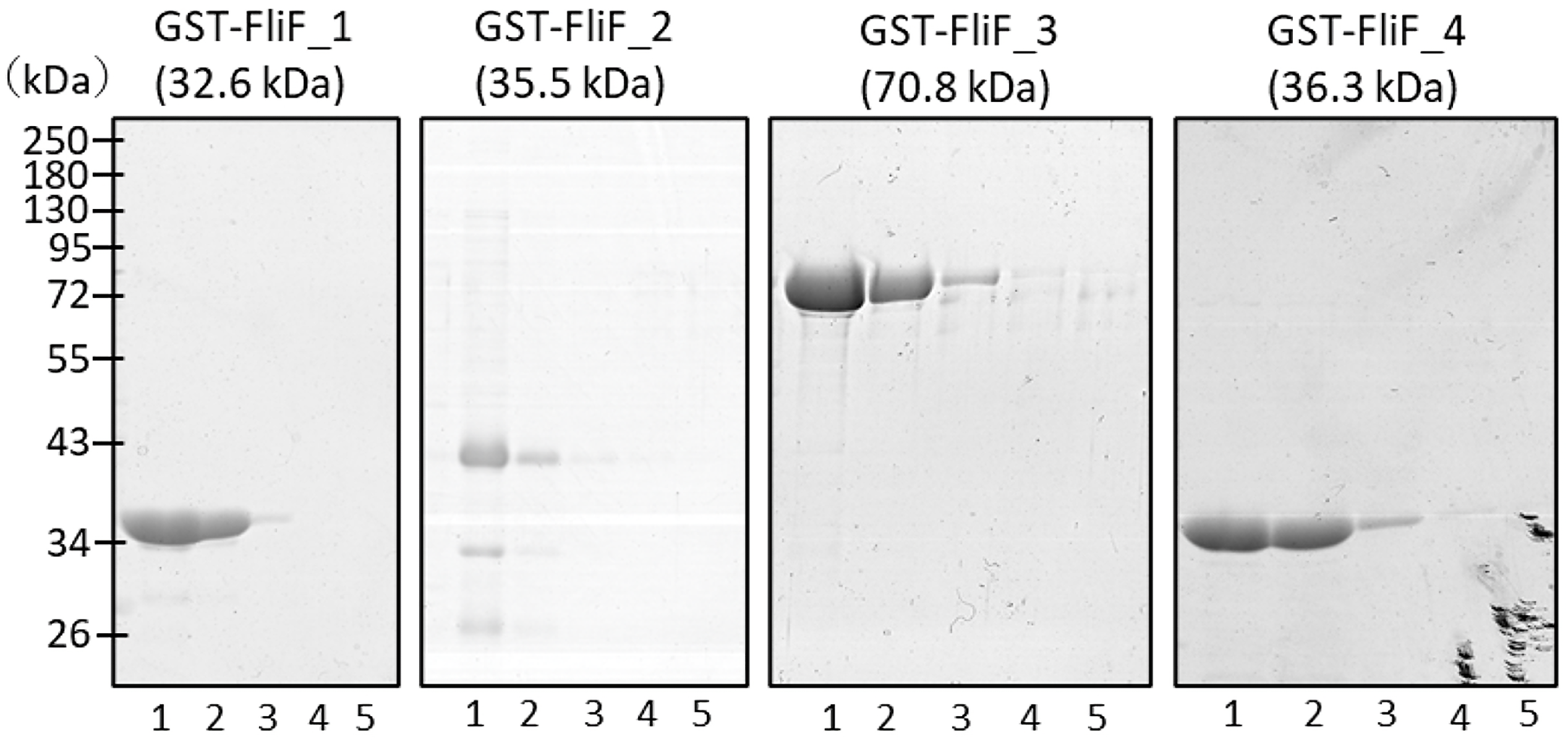
Purification of the FliF fragments. The FliF fragments, GST-FliF_1, GST-FliF_2, GST-FliF_3, and GST-FliF_4 were expressed from the plasmids, pYF101, pYF102, pYF103, and pYF104, respectively, in *E. coli* BL21(DE3) and purified using a GSTrap column. Eluted fractions of one to five of each fragment from the GSTrap column were separated by SDS-PAGE and subjected to CBB staining. The expected molecular weight of GST-fused FliF is shown in parentheses.

### Interaction detection between FlhF and purified FliF fragments

Binding experiments were performed using purified ST-FliF_1, GST-FliF_2, GST-FliF_3, GST-FliF_4, and cell lysate containing FlhF. FlhF was overproduced in *E. coli* BL21 (DE3) cells carrying the plasmid pTSK110, as previously described (22). After mixing with the purified GST-FliF fragment and cell lysate containing overproduced FlhF, Glutathione Sepharose 4B was added to the solution and allowed to react for 10 min at room temperature. GST-fused FliF fragments were purified and detected as expected; however, no specific binding of FlhF to the FliF fragments was observed.

### Detection of interaction by co-expression of FlhF and FliF

The binding experiment, which involved mixing the purified proteins and FlhF containing cell lysates directly, failed to detect an interaction between FliF and FlhF. FlhF belongs to the same family of signal recognition particles/receptors. It is a member of the Ffh/FtsY family, which may interact with the nascent chain FliF in the export pathway of membrane protein transport. Therefore, we examined binding by co-expressing FlhF and GST-FliF in the same *E. coli* cells. For this purpose, we transformed the plasmids pAK721 (express both FlhF and FlhG) into cells containing the plasmid that produces a FliF fragment. Co-expression of FlhF and FlhG somehow stabilized FlhF protein and more FlhF can be obtained in the soluble fraction, so that we used pAK721 for the binding experiment. *E. coli* cells were cultured to induce the expression of FlhF (and FlhG) and GST-FliF fragments. The soluble fraction was loaded into a 1 ml GST column and eluted. As shown in Fig. 3 and Fig. S3, FlhF bands were detected upon co-expression with GST-FliF_2 and GST-FliF_3. These results suggest that the first transmembrane region in the periplasmic region of FliF interacts with FlhF. We also used plasmid pAK322 that expresses FlhF only, instead of pAK721, for the same binding experiments. The results were essentially the same: FlhF can be co-eluted with GST-FliF_2 or GST-FliF_3. Therefore, co-expression of FlhG did not affect the FliF-FlhF binding. It is noteworthy that while purifying GST-FliF_2, two additional bands were observed at approximately 30 kDa and 26 kDa, excluding FlhF (Fig. S4). We used mass spectrometry to try to identify these proteins, and they appeared to be derived from β-lactamase.

**FIG 3.**
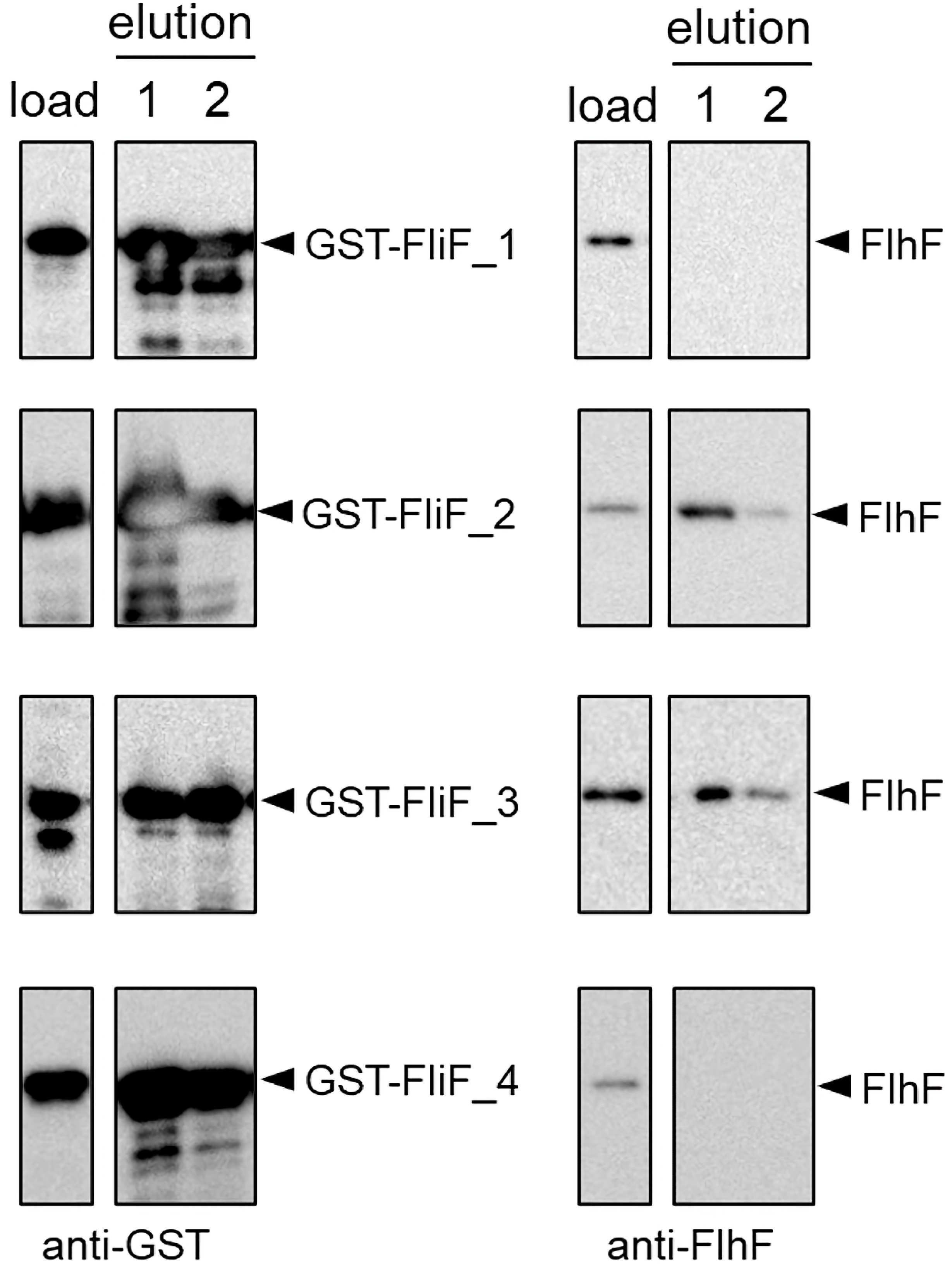
The binding experiment of the FliF fragments against FlhF. FliF fragments and FlhF were co-expressed in *E. coli* cells. The cytoplasmic fractions (load) were applied to a GSTrap column and eluted by glutathione. The first two eluted fractions (elution) were subjected to SDS-PAGE and Western blotting using an anti-FlhF antibody (α-FlhF) or anti-GST antibody (α-GST).

### Specifying the periplasmic region of the FliF interaction site against FlhF

To confirm which part of the periplasmic region of FliF is responsible for interaction with FlhF, we constructed three novel FliF fragments. These include FliF (88–469), FliF (109–469), and FliF (131–469) by slightly shaving the N-terminal side of the FliF_3 construct, 74–469 (Fig. 4A). Co-expression experiments were performed as described above. FliF (88–469) still retained its ability to bind FlhF, but FliF (109–469) and FliF (131–469) lost this ability (Fig. 4B). In the periplasmic region, there appears to be a binding site at 35 residues from 74 to 108 on the N-terminal side. Our analysis of the amino acid sequence of the region demonstrated numerous hydrophobic amino acids in approximately 20 residues of the N-terminal region of the fragment. Therefore, we speculate that this hydrophobic region is critical for the interaction between FlhF and FliF.

**FIG 4.**
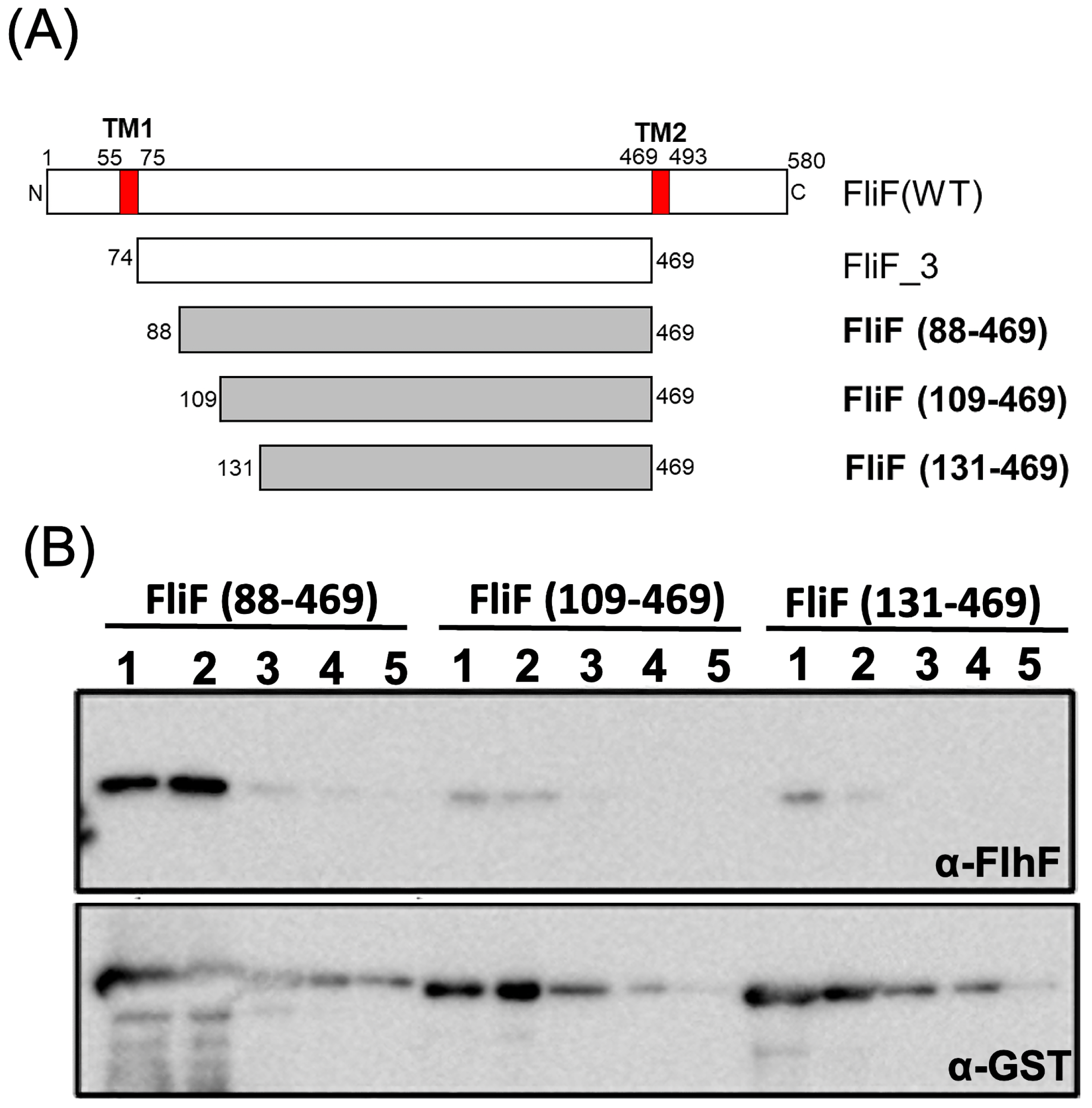
Narrowing the interaction site of the periplasmic region of FliF against FlhF. (A) Schematic diagram of the FliF fragments formed by truncating the N-terminal side of FliF_3 of pYF103. The deletions of the residues from 74 to 87, 108, and 130 were constructed and the fragments of FliG were named FliF (88–469), FliF (109–469), and FliF (181–469), respectively. (B) Western blotting to detect FlhF. Fractions of one to five eluted from the GSTrap column were separated by SDS-PAGE and proteins were detected using an anti-FlhF antibody (α-FlhF) or anti-GST antibody (α-GST).

## DISCUSSION

The assembly of the large MS-ring complex in the flagellar basal body remains uncertain. As described in the introduction, the MS-ring initiates flagellar morphogenesis. The complex structure of the MS-ring is composed of only one type of protein, FliF. FliF is a transmembrane (TM) protein with a large periplasmic region flanked by the N-terminal and C-terminal TM regions (Fig. 1 and Fig. S1). To assemble the MS-ring, FliF should be properly inserted into the membrane. Membrane proteins are usually inserted into the Sec system (23, 24). FlhF may recognize nascent FliF, which is transported to the Sec translocon in the membrane. Furthermore, SRP binds to sites with hydrophobic residues, suggesting that FlhF recognizes the TM1 region of FliF. In this study, it was demonstrated that approximately 20 residues from the N-terminal end of the first transmembrane region to the periplasmic region of FliF interact with FlhF. As Ffh recognizes and binds to the signal sequences carried by secreted proteins, FtsY transports Ffh-bound nascent polypeptides to the protein export machinery at the plasma membrane, where they are inserted. Based on our findings, FliF contains approximately 20 residues at the N-terminal end of the first transmembrane to the periplasmic region, which is among the 32 residues that serve as signal sequences. We found numerous hydrophobic amino acids on the N-terminal side of the periplasmic region of FliF (74–98). This region corresponds to the RBM1 region, which has been shown to have two α-helix structures in *Salmonella*. We speculate that this region may play a role in interacting with FlhF and assisting in the interaction with the translocon of the *E. coli* Sec system.

We have previously examined the roles of the N- and C-terminal regions of FliF (25). When MS rings were purified from *E. coli* cells that produce ΔC83FliF (which comprises a C-terminal deletion of 83 cytoplasmic residues of FliF and is not functional in *Vibrio* cells), we could recover more MS rings with FlhF than without FlhF as wild-type FliF. When MS rings purified from *Vibrio* cells derived from *E. coli* cells producing ΔN30FliF or ΔN50FliF (the N-terminal deletions of the cytoplasmic 30 and 50 residues of FliF) are compared with MS rings purified from wild-type FliF, fewer MS rings are formed. As a result, we concluded that the N-terminal region of FliF is necessary for the FlhF function to assemble FliF. Interaction of FlhF with only the N-terminal cytoplasmic region of FliF (FliF_1) was not detected in this study. We believe that the essential region of FliF for interaction with FlhF is the TM1 region. Additionally, the hydrophobic N-terminal periplasmic region of FliF and the N-terminal cytoplasmic region of FliF may facilitate the interaction of FlhF with the FliF signal sequence, contributing to MS-ring stability or ring formation after membrane insertion.

GST pull-down experiments were used in this study to identify the region of FliF that interacts with FlhF. However, it remains unclear how FlhF contributes to MS-ring formation. In the future, we will conduct further FlhF-FliF-binding experiments using *Salmonella* FliF. Additionally, *Salmonella* FliF has a sequence similar to that of *V. alginolyticus*. In contrast, *Salmonella* lacks FlhF, and FliF is not localized to the cell pole. These results suggest that FlhF binds specifically to *Vibrio* FliF or non-specifically to the hydrophobic region, and we are investigating FlhF-FliF binding using *flhF* and *fliF* mutants.

## MATERIALS AND METHODS

### Strains, plasmids, and culture conditions

The strains and plasmids used are listed in Table S1. *Escherichia coli* was cultured in LB medium [1% (w/v) Bacto Tryptone, 0.5% (w/v) yeast extract, and 0.5% (w/v) NaCl] at 37°C. To create a solid medium, agar was added at 1.25% (w/v) to the LB medium. If necessary, chloramphenicol (Cm) and ampicillin (Amp) were added at final concentrations of 25 μg/mL and 100 μg/mL, respectively. The gene under the control of the arabinose promoter (*pBAD*) was induced by adding an amount of arabinose (0.02%) suitable for the expression level.

### DNA manipulations, mutagenesis and sequencing

Routine DNA manipulations were performed according to standard procedures. To produce FliF fragments with N-terminal fusion of GST, the *fliF* fragments were amplified by PCR using PfuUltra HF DNA Polymerase using the forward and reverse primers designated for FliF_1, FliF_2, FliF_3, and FliF_4 (Fig. 1). To fuse GST at the restriction enzyme site, the fragments were cloned into the vector plasmid pGEX-6p-2 at BamHI and XhoI sites.

Further N-terminal deletions, FliF (88–469), FliF (109–469), and FliF (131–469), were constructed by cutting the N-terminal residues of FliF_3 and FliF (74–469) of pYF103 using the inverse PCR method as previously described (22).

### Purification of GST-FliF fragments

The *E. coli* BL21(DE3) cells were transformed with pYF101, pYF102, pYF103, or pYF104 vectors and stored in a freezer at −80 °C. Each strain was inoculated into an LB medium containing ampicillin, and on the following day, 0.35 mL of the overnight culture was inoculated into 35 mL of LB medium containing ampicillin. Incubation was maintained for 3 h at 37°C after OD_660_ reached 0.5 and IPTG was added at a final concentration of 0.5 mM. Cells were collected by low-speed centrifugation, suspended in 25 mL of PBS (137 mM NaCl, 2.68 mM KCl, 10 mM Na_2_HPO_4_, 2 mM KH_2_PO_4_), and sonicated (large probe, power = 8, duty cycle = 50%, 50 seconds, three times). Unbroken cells were removed using low-speed centrifugation, and the supernatant was recovered by high-speed centrifugation (150000 *× g*, 30 min, 4 °C). The cytoplasmic fraction (supernatant after ultracentrifugation) was loaded onto a GSTrap (FF) 1 ml column (GST) using a peristaltic pump. After washing the column with PBS, elution buffer (50 mM Tris-HCl, pH 8, 10 mM reduced glutathione) was added to the column using a syringe to collect 1 mL for each elution. Purified proteins were frozen in liquid nitrogen and stored at −80 °C. The protein concentration was calculated from the absorbance at wavelengths of 280 nm and 320 nm measured using a spectrophotometer.

### Interaction between purified GST-FliF fragments and FlhF

FlhF was obtained as previously described (22). *Escherichia coli* BL21 (DE3) cells expressing pTSK110 were grown in an LB medium containing ampicillin at 37°C. At 0.5 OD_660,_ FlhF protein was induced by cold shock (on ice for 5 min) and IPTG, then incubated at 16°C overnight. The cells were collected by centrifugation and suspended in FlhF buffer [20 mM Tris-HCl (pH 8.0), 300 mM NaCl, 10 mM MgCl_2_, and 10 mM KCl] containing a complete protease inhibitor (Roche) and then sonicated (power = 5, duty cycle = 50%, 30 s, five times). The unbroken cells were removed by low-speed centrifugation, and the supernatant was ultracentrifuged (154,000 *×g*, 30 min, 4°C). The resulting supernatant was mixed with glutathione sepharose 4 B, which had been bound to GST-fused FliF. After washing with PBS, the bound proteins were eluted with the elution buffer.

### Binding detection between GST-FliF fragments and FlhF in cells

*Escherichia coli* BL21 (DE3) cells harboring pYF101, pYF102, pYF103, pYF104 or pYF103 derivatives (pYF1031, pYF1032, pYF1033) were transformed with pAK721. The cells containing two plasmids were grown in LB medium containing Amp and Cm at 37°C; 0.2% arabinose was added at the start of culture, and IPTG was added at 0.5 of OD_660_. The cells were further incubated at 18°C for 3 h, collected by centrifugation, and suspended in FlhF buffer; then the cells were sonicated. The unbroken cells were removed by low-speed centrifugation, and the supernatant was ultracentrifuged (154,000 *×g*, 30 min, 4°C). The resulting supernatant, that contains both FlhF and GST-FliF fragments, was loaded onto a GSTrap (FF) 1 ml column using a peristaltic pump. After washing the column with PBS, elution buffer (50 mM Tris-HCl, pH 8, 10 mM reduced glutathione) was added to the column using a syringe to collect 1 mL for each elution.

## ACKNOWLEDGEMENTS

This work was supported, in part, by JSPS KAKENHI: Grant Number; 20H03220 (to M.H.).

## REFERENCES

1. Laloux G, Jacobs-Wagner C. 2014. How do bacteria localize proteins to the cell pole? J Cell Sci 127:11–9.

2. Keilberg D, Søgaard-Andersen L. 2014. Regulation of bacterial cell polarity by small GTPases. Biochemistry 53:1899–907.

3. Kojima S, Terashima H, Homma M. 2020. Regulation of the single polar flagellar biogenesis. Biomolecules 10:533.

4. Kusumoto A, Kamisaka K, Yakushi T, Terashima H, Shinohara A, Homma M. 2006. Regulation of polar flagellar number by the flhF and flhG genes in Vibrio alginolyticus. J Biochem 139:113–21.

5. Bange G, Sinning I. 2013. SIMIBI twins in protein targeting and localization. Nat Struct Mol Biol 20:776–80.

6. Steinberg R, Knupffer L, Origi A, Asti R, Koch HG. 2018. Co-translational protein targeting in bacteria. FEMS Microbiol Lett 365:fny095.

7. Kusumoto A, Shinohara A, Terashima H, Kojima S, Yakushi T, Homma M. 2008. Collaboration of FlhF and FlhG to regulate polar-flagella number and localization in Vibrio alginolyticus. Microbiology (Reading) 154:1390–1399.

8. Yamaichi Y, Bruckner R, Ringgaard S, Moll A, Cameron DE, Briegel A, Jensen GJ, Davis BM, Waldor MK. 2012. A multidomain hub anchors the chromosome segregation and chemotactic machinery to the bacterial pole. Genes Dev 26:2348–60.

9. Takekawa N, Kwon S, Nishioka N, Kojima S, Homma M. 2016. HubP, a Polar Landmark Protein, Regulates Flagellar Number by Assisting in the Proper Polar Localization of FlhG in Vibrio alginolyticus. J Bacteriol 198:3091–3098.

10. Inaba S, Nishigaki T, Takekawa N, Kojima S, Homma M. 2017. Localization and domain characterization of the SflA regulator of flagellar formation in Vibrio alginolyticus. Genes Cells 22:619–627.

11. Kitaoka M, Nishigaki T, Ihara K, Nishioka N, Kojima S, Homma M. 2013. A novel dnaJ family gene, sflA, encodes an inhibitor of flagellation in marine Vibrio species. J Bacteriol 195:816–22.

12. Sakuma M, Nishikawa S, Inaba S, Nishigaki T, Kojima S, Homma M, Imada K. 2019. Structure of the periplasmic domain of SflA involved in spatial regulation of the flagellar biogenesis of Vibrio reveals a TPR/SLR-like fold. J Biochem 166:197–204.

13. Homma M, Takekawa N, Fujiwara K, Hao Y, Onoue Y, Kojima S. 2022. Formation of multiple flagella caused by a mutation of the flagellar rotor protein FliM in Vibrio alginolyticus. Genes Cells 27:568–578.

14. Homma M, Kobayakawa T, Hao Y, Nishikino T, Kojima S. 2022. Function and structure of FlaK, a master regulator of the polar flagellar genes in marine Vibrio. J Bacteriol 204:e0032022.

15. Homma M, Aizawa S-I, Dean GE, Macnab RM. 1987. Identification of the M-ring protein of the flagellar motor of Salmonella typhimurium. Proc Natl Acad Sci USA 84:7483–7487.

16. Ueno T, Oosawa K, Aizawa S. 1994. Domain structures of the MS ring component protein (FliF) of the flagellar basal body of Salmonella typhimurium. J Mol Biol 236:546–555.

17. Kawamoto A, Miyata T, Makino F, Kinoshita M, Minamino T, Imada K, Kato T, Namba K. 2021. Native flagellar MS ring is formed by 34 subunits with 23-fold and 11-fold subsymmetries. Nat Commun 12:4223.

18. Takekawa N, Kawamoto A, Miyata T, Sakuma M, Kato T, Kojima S, Kinoshita M, Minamino T, Namba K, Homma M, Imada K. 2021. Two distinct conformations in 34 FliF subunits generate three different symmetries within the flagellar MS-ring. MBio 12:e03199–20.

19. Johnson S, Fong YH, Deme JC, Furlong EJ, Kuhlen L, Lea SM. 2020. Symmetry mismatch in the MS-ring of the bacterial flagellar rotor explains the structural coordination of secretion and rotation. Nat Microbiol 5:966–975.

20. Ogawa R, Abe-Yoshizumi R, Kishi T, Homma M, Kojima S. 2015. Interaction of the C-terminal tail of FliF with FliG from the Na^+^-driven flagellar motor of Vibrio alginolyticus. J Bacteriol 197:63–72.

21. Terashima H, Hirano K, Inoue Y, Tokano T, Kawamoto A, Kato T, Yamaguchi E, Namba K, Uchihashi T, Kojima S, Homma M. 2020. Assembly mechanism of a supramolecular MS-ring complex to initiate bacterial flagellar biogenesis in Vibrio species. J Bacteriol 202:e00236.

22. Kondo S, Imura Y, Mizuno A, Homma M, Kojima S. 2018. Biochemical analysis of GTPase FlhF which controls the number and position of flagellar formation in marine Vibrio. Sci Rep 8:12115.

23. Smets D, Loos MS, Karamanou S, Economou A. 2019. Protein transport across the bacterial plasma membrane by the Sec pathway. Protein J 38:262–273.

24. Cranford-Smith T, Huber D. 2018. The way is the goal: how SecA transports proteins across the cytoplasmic membrane in bacteria. FEMS Microbiol Lett 365:fny093.

25. Kojima S, Kajino H, Hirano K, Inoue Y, Terashima H, Homma M. 2021. Role of the N- and C-terminal regions of FliF, the MS ring component in Vibrio flagellar basal body. J Bacteriol 203:e00009–21.

